# Biologics-associated Risks for Incident Skin and Soft Tissue Infections in Psoriasis Patients: Results from Propensity Score-Stratified Survival Analysis

**DOI:** 10.1101/126383

**Authors:** Su-Hsun Liu, Tsung-Han Hsieh, Leslie Y Chen, Yhu-Chering Huang, Yu-Huei Huang

## Abstract

**Background:** How biologics affect psoriasis patients’ risks for SSTIs in a pragmatic clinical setting remains unclear.

**Methods:** In a cohort of adult psoriasis outpatients (aged 20 years or older) who visited the Dermatology Clinic in 2010-2015, we compared incident SSTI risks between patients using biologics (users) versus nonbiologics (nonusers). We also estimated SSTI risks in biologics-associated time-periods relative to nonbiologics only in users. We applied random effects Cox proportional hazard models with propensity score-stratification to account for differential baseline hazards.

**Results:** Over a median follow-up of 2.8 years (interquartile range: 1.5, 4.3), 172 of 922 patients ever received biologics (18.7%); 233 SSTI incidents occurred during 2518.3 person-years, with an overall incidence of 9.3/100 person-years (95% confidence interval [CI]: 8.1, 10.6). In univariate analysis, users showed an 89% lower risk for SSTIs than nonusers (hazard ratio [HR]: 0.11, 95%CI: 0.05, 0.26); the association persisted in a multivariable model (adjusted HR: 0.26, 95%CI: 0.12, 0.58). Among biologics users, biologics-exposed time-periods were associated with a nonsignificant 21% increased risk (adjusted HR: 1.21, 95%CI: 0.41, 3.59).

**Conclusions:** Despite of adjusting for the underlying risk profiles, risk comparisons between biologics users and nonusers remained confounded by treatment selection. By comparing time-periods being exposed versus unexposed to biologics among users, the current analysis did not find evidence for an increased SSTI risk that was associated with biologics use in psoriasis patients.

## BACKGROUND

*Staphylococcus aureus* (*S. aureus*) remains a major cause of skin and soft tissue infections (SSTI) worldwide,[1] particularly by the resistant strains-methicillin-resistant *S. aureus* (MRSA).[2] The emergence of community-associated MRSA further raises concerns about hidden reservoirs of asymptomatic carriage in the community setting.[3] While as many as 20-50% of healthy adults in the general population may carry *S. aureus*,[3] patients with atopic dermatitis (AD) or psoriasis reportedly share a common predilection for *S. aureus* colonisation.[4-6]

With the accumulating success in treating chronic immune-mediated diseases including psoriasis, biologic agents have appeared safe in a trial setting.[7, 8] The most common infections reported by controlled trials and their long-term extension studies, however, were upper respiratory tract infections.[7-9] Cutaneous infections and infestations seemed uncommon among psoriasis patients treated with biologics (4.4/100 person-years) as compared to those using classic systemic drugs (4.7/100 person-years) even with an extended observation.[10]

Although such uncommonness of SSTI in psoriasis patients was consistent with early clinical observations,[11] such rarity has contradicted with recent findings that bacterial infection or colonization correlated with disease activities in psoriasis patients.[12-14] The discrepancy in the literature[8, 10, 12, 13, 15-24] may stem from different perspectives on inflamed skin lesions as a treatment-associated complication or a clinical manifestation of the disease per se; the distinction of which depends on swab cultures that are not part of the routine care.[25, 26] Differences in how comparators are selected in evaluating infection risks could also affect conclusions made. In controlled trials, comparison groups randomly receive treatment regimens so as to ensure comparability across groups; in practice, prescription of biologics is a nonrandom decision but guided in a hierarchical, stage-by-stage fashion.[26] Patients who are advanced to biologics therapies are clinically different from those who can benefit from conventional regimens not only in disease severity but also in sociodemographic and comorbidities.[26] While admitting this incomparability, few cohort studies utilizing data from patient registries[10, 20-22] healthcare claims data[23, 24] attended to this confounding-by-indication.

Therefore, in the current study, we sought to determine biologics-associated risks for SSTIs among psoriasis patients in a pragmatic clinical setting. Among possible alternatives,[27] we employed propensity scores to facilitate fair comparisons between patients who ever used (users) versus those who received only nonbiologics (nonusers) during the study period, by taking into account the predicted probability of receiving the exposure (or treatment) of interest.[28-30] In addition, we aimed to quantify and compare SSTI risks in time-periods exposed to biologics versus nonbiologics among biologics-users only.

## PATIENTS AND METHODS

### Study design and patient population

Using ICD-9-CM (International Classification of Diseases, Ninth Revision, Clinical Modification) codes of 696.0 and 696.1, we retrospectively identified a cohort of psoriasis patients in the Electronic Medical Database (EMD). We included only adults (aged 20 years or older) who visited the Dermatology Outpatient Clinic at least twice within a moving 365-day window between January 1st, 2010 and August 31st, 2015, with the latter being the administrative censoring date. The Institutional Review Board of Chang Gung Medical Institution reviewed and approved the study protocol and the analytic plan; the Institutional Review Board also waived the requirement for obtaining consent forms.

### Data collection

#### Exposures

We classified patients’ prescription medications into either biologic (including Adalimumab, Etanercept, Golimumab, or Ustekinumab) or nonbiologics at each clinical encounter. Based on the reported bioavailability half-life of each biologic in the individual pharmaceutical pamphlets (Adalimumab: 20 days; Etanercept: 5 days; Golimumab: 14 days; Ustekinumab: 21 days), we assumed a maximum person-time being exposed to each biologic at the censoring visit. However, we did not assume any lagged effect of biologics on patients’ risk for an incident SSTI.

#### Outcomes

We also used ICD-9-CM codes to identify SSTIs in patients’ medical records as previously reported.[31] When there were more than one SSTI-related code at the same visit, we favoured a more specific diagnosis than a less specific one; for example, surgical site infections (SSI, 998.5x or 999.3x) was chosen over nonspecified infections (686.x).

#### Covariates

We collected patients’ demographic characteristics, including age at study entry; sex; laboratory data, including liver function tests (AST, aspartate transaminase; ALT, alanine transaminase), lipid profile (total cholesterol; LDL, low density lipoprotein-cholesterol; triglyceride), fasting or post-prandial plasma glucose level, glycohemoglobulin, and hepatitis B and C profile; blood or tissue culture results during the same observation period whenever available. Additionally, we quantified patients’ use of corticosteroids by converting the average daily dose of glucocorticoid-containing products to equivalents of 5mg-prednisolone at study entry and follow-up using a free online converting algorithm.[23, 32] We categorized patients using systemic (oral or intra-muscular) glucocorticoids into a low- (<5 mg) or medium-to-high (≥ 5 mg) group; topical glucocorticoids only; or nonsteroid-containing medications. We categorized antibiotics as prophylactic use when its prescription was not accompanied with a new SSTI-associated diagnostic code at the same visit.

## Statistical analysis

We described and compared patient characteristics at study entry and follow-up between patients ever and never receiving biologics during the study period. We calculated event rates and associated 95% confidence intervals [CI] for the first incident SSTI by patients’ characteristics. We constructed Gaussian-distributed random effects Cox proportional hazards models to account for repeated visits per subject.[33] Each patient entered a risk set for an incident when s/he entered the study without a prevalent SSTI or when a prevalent SSTI episode had cleared. Censoring occurred at the time of the outcome; when the patient no longer returned to the clinic or was administratively censored. We calculated a mid-interval date between two consecutive visits without and with an SSTI-associated diagnosis as an event time.[34]

Propensity scores were predicted probability estimates of a logistic regression model,[35] that included patients’ sex and clinical characteristics at study entry, such as age, calendar year, and the initial presence of psoriatic arthropathy or not. Similar to using propensity score matching,[23] the stratification method in survival analysis allowed for varying ‘baseline’ hazards across the propensity score-based strata (quintile bins) while the inclusion of only pre-treatment covariates avoided biases that might be introduced by post-treatment covariates.[28, 36] We performed descriptive analysis and calculated propensity scores in Stata (version 13.0);[37] constructed Cox regression models in R (version 3.3.1).[38, 39] We replied on Akaike information criteria (AIC) that were derived from the penalized log-likelihood function[39] to guide model comparisons. All analyses were at two-tailed significance level of 0.05.

## RESULTS

Among 1,672 psoriasis outpatients (29,575 visits) identified in 2010-2015, 40 patients were younger than 20 years and thus excluded (540 visits); 11 patients entered the study at ages 17-19 years and we retained only their adult records (136 visits) in the following analysis. With additional exclusion of 764 patients (Figure 1), 959 adult patients met our inclusion criteria, 37 of whom had a prevalent SSTI-associated diagnosis at the study entry (3.9%) and were further excluded (Supplementary Table S1).

**Figure 1.**
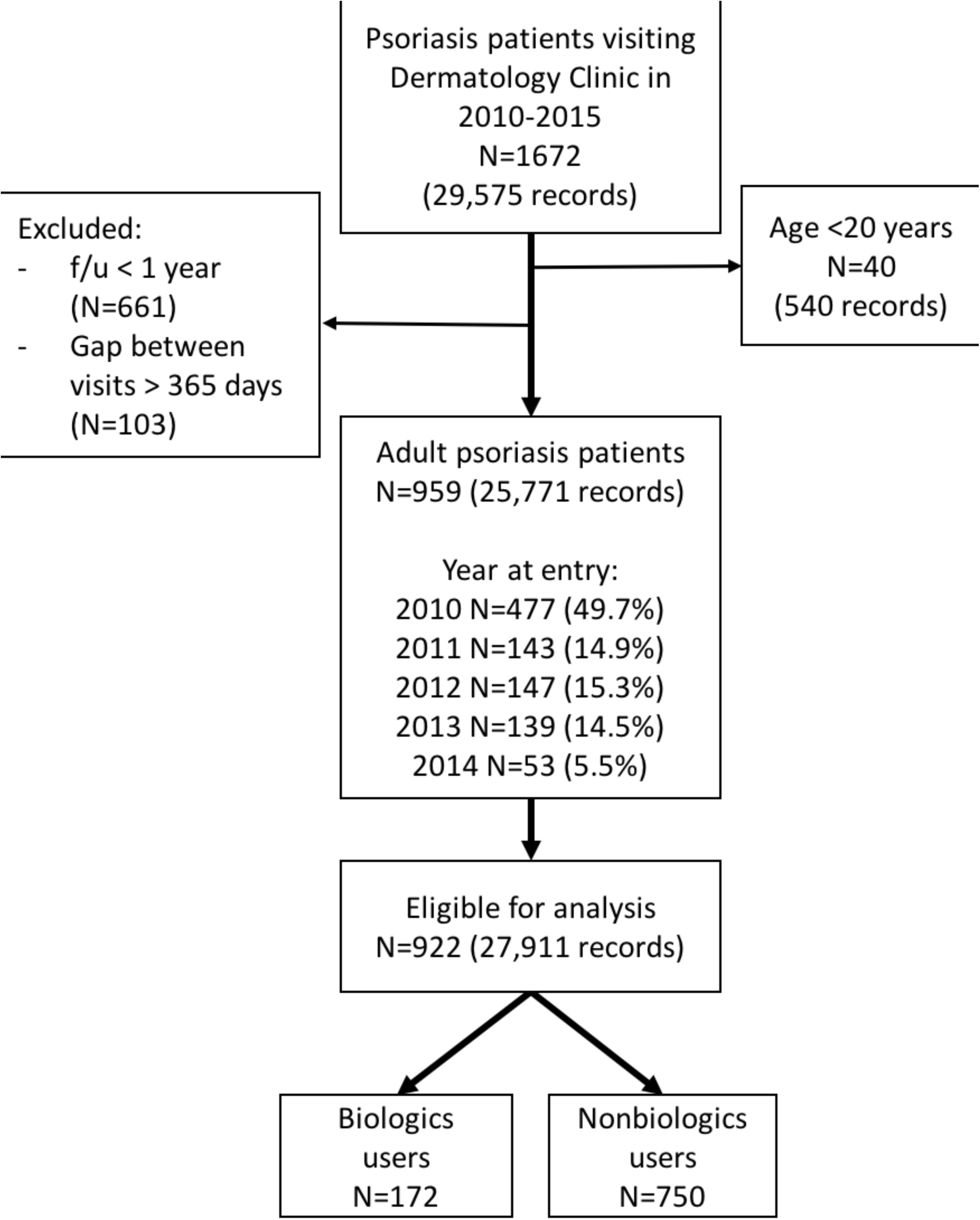
Flowchart of the identification and selection procedure for eligible psoriasis patients included in the analysis.

Overall, we identified 922 psoriasis patients among whom 172 ever received biologics treatment (14.6%, Table 1). More than 12% of the study population was 65 years or older (12.8%); 70.4% were men; 14.1% also had a diagnosis of psoriatic arthropathy and 76.2% were using topical steroids at the first eligible visit. On average, biologics users were 6.5 years younger (median: 43.5 vs. 50 years, P < 0.001) and more likely to have join involvement than nonusers (42.0% vs. 11.7%, P < 0.001) at the study entry. As compared to nonusers, more users ever received systematic glucocorticoids (21.3% vs. 5.5%) at follow-up and were more likely to have multiple metabolic risks factors, including elevated liver enzymes, hyperlipidemia, and impaired glucose metabolism or a history of diabetes (Table 1). Prophylactic antibiotics were infrequent, with an overall period prevalence of 0.6% (N=8) during a median follow-up period of 2.8 years (interquartile range [IQR]: 1.5-4.3).

**Table 1.**
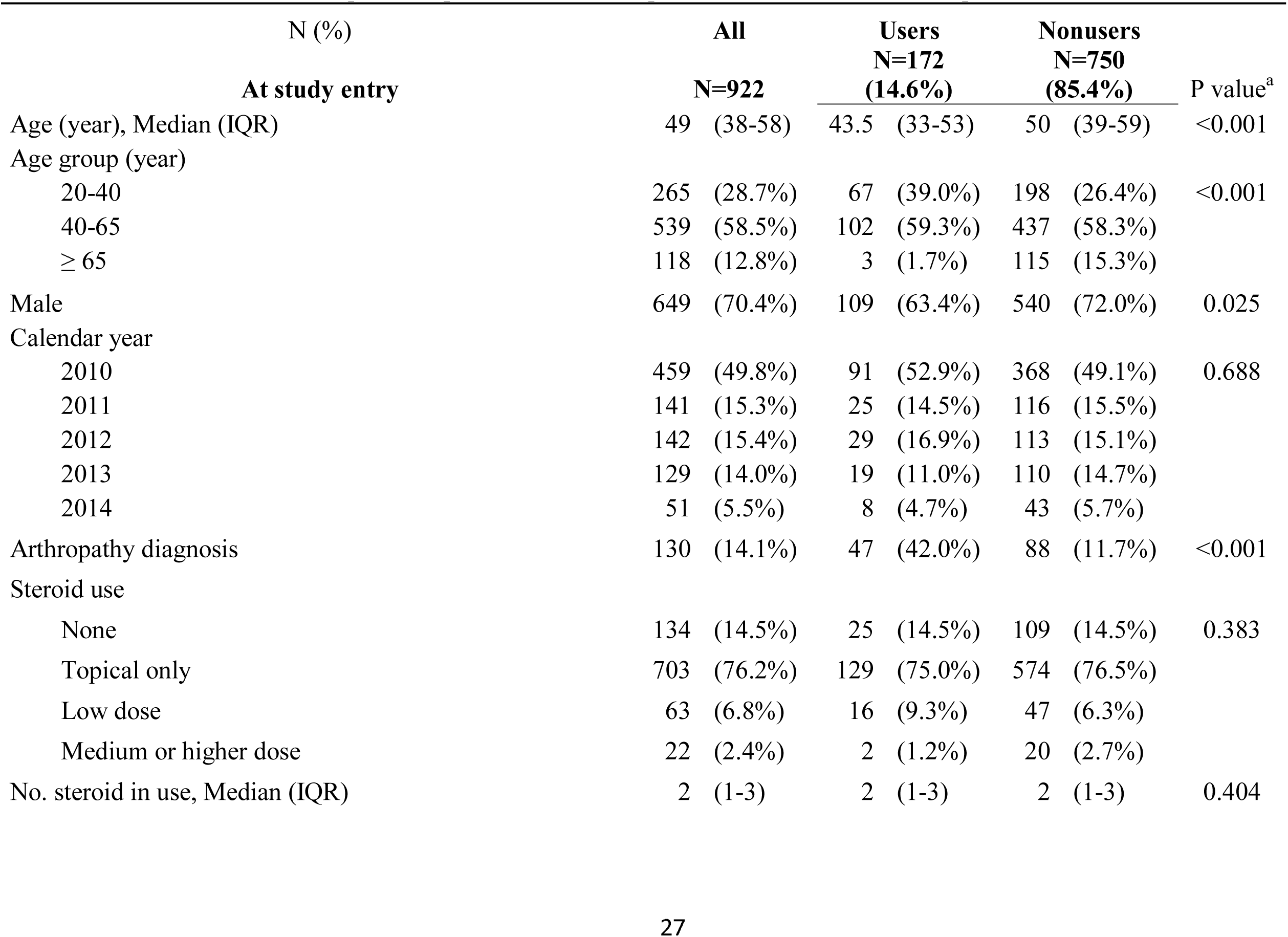
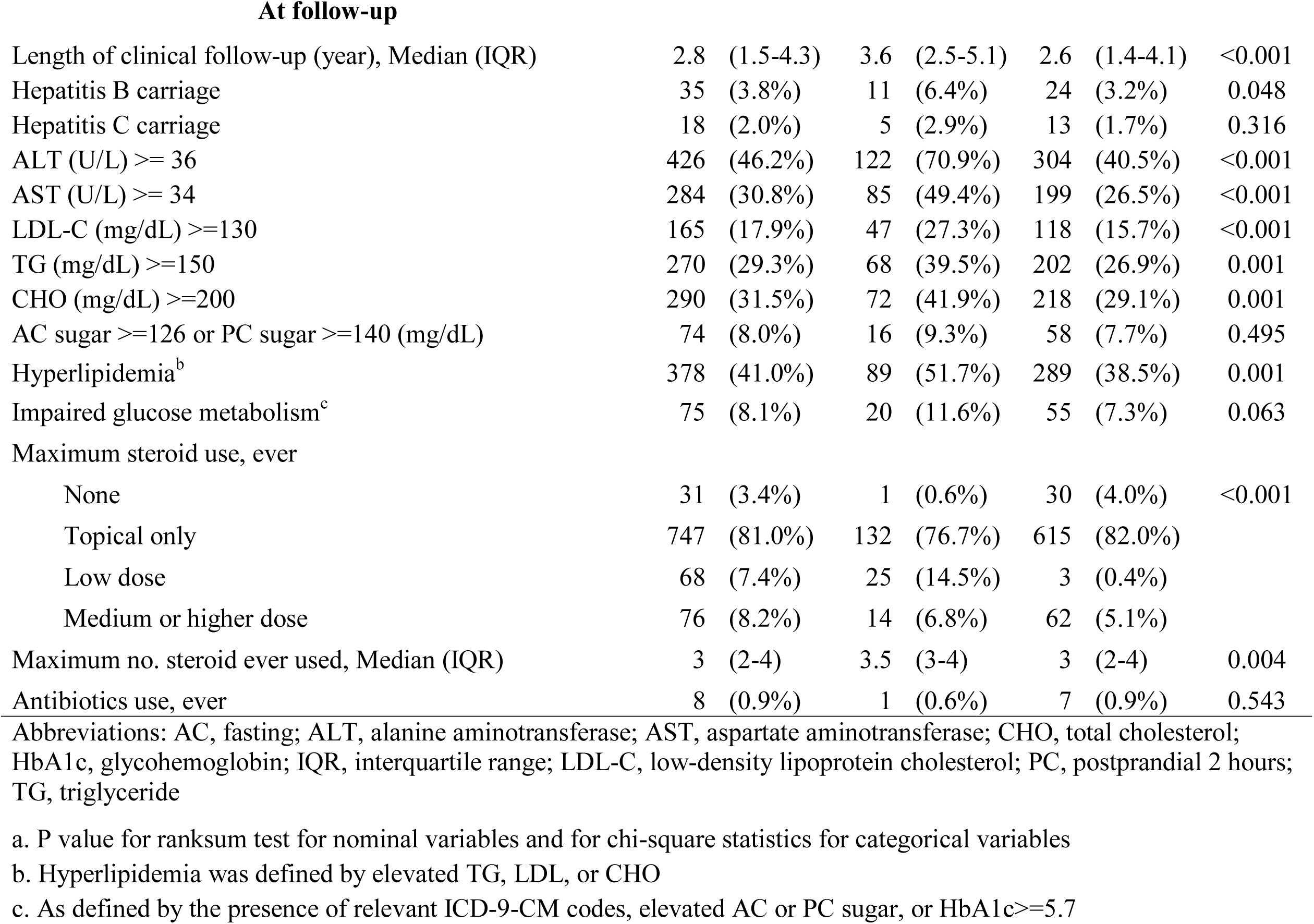
Characteristics of adult psoriasis patients who developed incident SSTIs at follow-up in 2010-2015

|  | N (%) | All<br>N=922 | Users<br>N=172<br>(14.6%) | Nonusers<br>N=750<br>(85.4%) | P value<sup>a</sup> |
| --- | --- | --- | --- | --- | --- |
| **At study entry** |  |  |  |  |  |
| Age (year), Median (IQR) |  | 49 (38-58) | 43.5 (33-53) | 50 (39-59) | <0.001 |
| Age group (year) |  |  |  |  |  |
| 20-40 |  | 265 (28.7%) | 67 (39.0%) | 198 (26.4%) | <0.001 |
| 40-65 |  | 539 (58.5%) | 102 (59.3%) | 437 (58.3%) |  |
| ≥ 65 |  | 118 (12.8%) | 3 (1.7%) | 115 (15.3%) |  |
| Male |  | 649 (70.4%) | 109 (63.4%) | 540 (72.0%) | 0.025 |
| Calendar year |  |  |  |  |  |
| 2010 |  | 459 (49.8%) | 91 (52.9%) | 368 (49.1%) | 0.688 |
| 2011 |  | 141 (15.3%) | 25 (14.5%) | 116 (15.5%) |  |
| 2012 |  | 142 (15.4%) | 29 (16.9%) | 113 (15.1%) |  |
| 2013 |  | 129 (14.0%) | 19 (11.0%) | 110 (14.7%) |  |
| 2014 |  | 51 (5.5%) | 8 (4.7%) | 43 (5.7%) |  |
| Arthropathy diagnosis |  | 130 (14.1%) | 47 (42.0%) | 88 (11.7%) | <0.001 |
| Steroid use |  |  |  |  |  |
| None |  | 134 (14.5%) | 25 (14.5%) | 109 (14.5%) | 0.383 |
| Topical only |  | 703 (76.2%) | 129 (75.0%) | 574 (76.5%) |  |
| Low dose |  | 63 (6.8%) | 16 (9.3%) | 47 (6.3%) |  |
| Medium or higher dose |  | 22 (2.4%) | 2 (1.2%) | 20 (2.7%) |  |
| No. steroid in use, Median (IQR) |  | 2 (1-3) | 2 (1-3) | 2 (1-3) | 0.404 |

At follow-up
|  |  |  |  |  |
| --- | --- | --- | --- | --- |
| Length of clinical follow-up (year), Median (IQR) | 2.8 (1.5-4.3) | 3.6 (2.5-5.1) | 2.6 (1.4-4.1) | <0.001 |
| Hepatitis B carriage | 35 (3.8%) | 11 (6.4%) | 24 (3.2%) | 0.048 |
| Hepatitis C carriage | 18 (2.0%) | 5 (2.9%) | 13 (1.7%) | 0.316 |
| ALT (U/L) $\geq 36$ | 426 (46.2%) | 122 (70.9%) | 304 (40.5%) | <0.001 |
| AST (U/L) $\geq 34$ | 284 (30.8%) | 85 (49.4%) | 199 (26.5%) | <0.001 |
| LDL-C (mg/dL) $\geq 130$ | 165 (17.9%) | 47 (27.3%) | 118 (15.7%) | <0.001 |
| TG (mg/dL) $\geq 150$ | 270 (29.3%) | 68 (39.5%) | 202 (26.9%) | 0.001 |
| CHO (mg/dL) $\geq 200$ | 290 (31.5%) | 72 (41.9%) | 218 (29.1%) | 0.001 |
| AC sugar $\geq 126$ or PC sugar $\geq 140$ (mg/dL) | 74 (8.0%) | 16 (9.3%) | 58 (7.7%) | 0.495 |
| Hyperlipidemia<sup>b</sup> | 378 (41.0%) | 89 (51.7%) | 289 (38.5%) | 0.001 |
| Impaired glucose metabolism<sup>c</sup> | 75 (8.1%) | 20 (11.6%) | 55 (7.3%) | 0.063 |
| Maximum steroid use, ever |  |  |  |  |
| None | 31 (3.4%) | 1 (0.6%) | 30 (4.0%) | <0.001 |
| Topical only | 747 (81.0%) | 132 (76.7%) | 615 (82.0%) |  |
| Low dose | 68 (7.4%) | 25 (14.5%) | 3 (0.4%) |  |
| Medium or higher dose | 76 (8.2%) | 14 (6.8%) | 62 (5.1%) |  |
| Maximum no. steroid ever used, Median (IQR) | 3 (2-4) | 3.5 (3-4) | 3 (2-4) | 0.004 |
| Antibiotics use, ever | 8 (0.9%) | 1 (0.6%) | 7 (0.9%) | 0.543 |

Abbreviations: AC, fasting; ALT, alanine aminotransferase; AST, aspartate aminotransferase; CHO, total cholesterol; HbA1c, glycohemoglobin; IQR, interquartile range; LDL-C, low-density lipoprotein cholesterol; PC, postprandial 2 hours; TG, triglyceride
a. P value for ranksum test for nominal variables and for chi-square statistics for categorical variables
b. Hyperlipidemia was defined by elevated TG, LDL, or CHO
c. As defined by the presence of relevant ICD-9-CM codes, elevated AC or PC sugar, or HbA1c $\geq 5.7$

During 2518.3 person-years of observation, we identified 233 incident SSTI events with an overall estimated incidence rate (IR) of 9.3/100 person-years (95%CI: 8.1, 10.6; Table 2). Biologics users (IR: 5.5/100 person-years, 95%CI: 4.0, 7.9) appeared to have a greatly reduced risk as compared to nonusers (IR: 10.4/100 person-years, 95%CI: 9.0, 12.1). A younger age (9.1/100 for aged 20-40 years vs. 11.2/100 person-years for aged 65 years or older) and an early entry into the study (8.8/100 in 2010 vs. 15.4/100 person-years in 2013) also appeared protective whereas hepatitis C carriage or antibiotic use was correlated with a higher SSTI rate (Table 2).

Before propensity score adjustment, biologics users also showed a significantly favourable survivorship from an incident SSTI comparing to nonusers (*P* for log-rank test: 0.002, Figure 2).

**Table 2.**
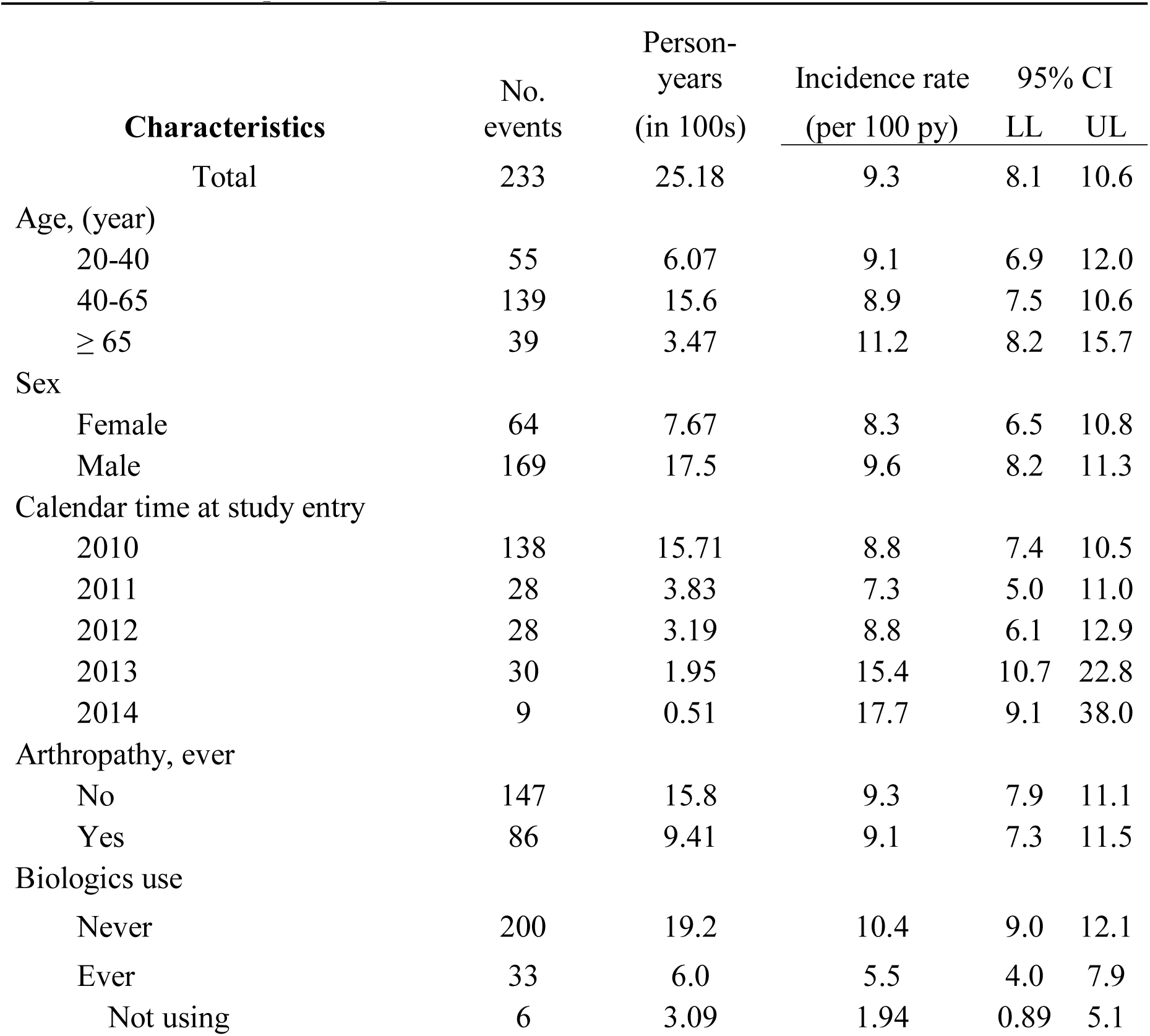
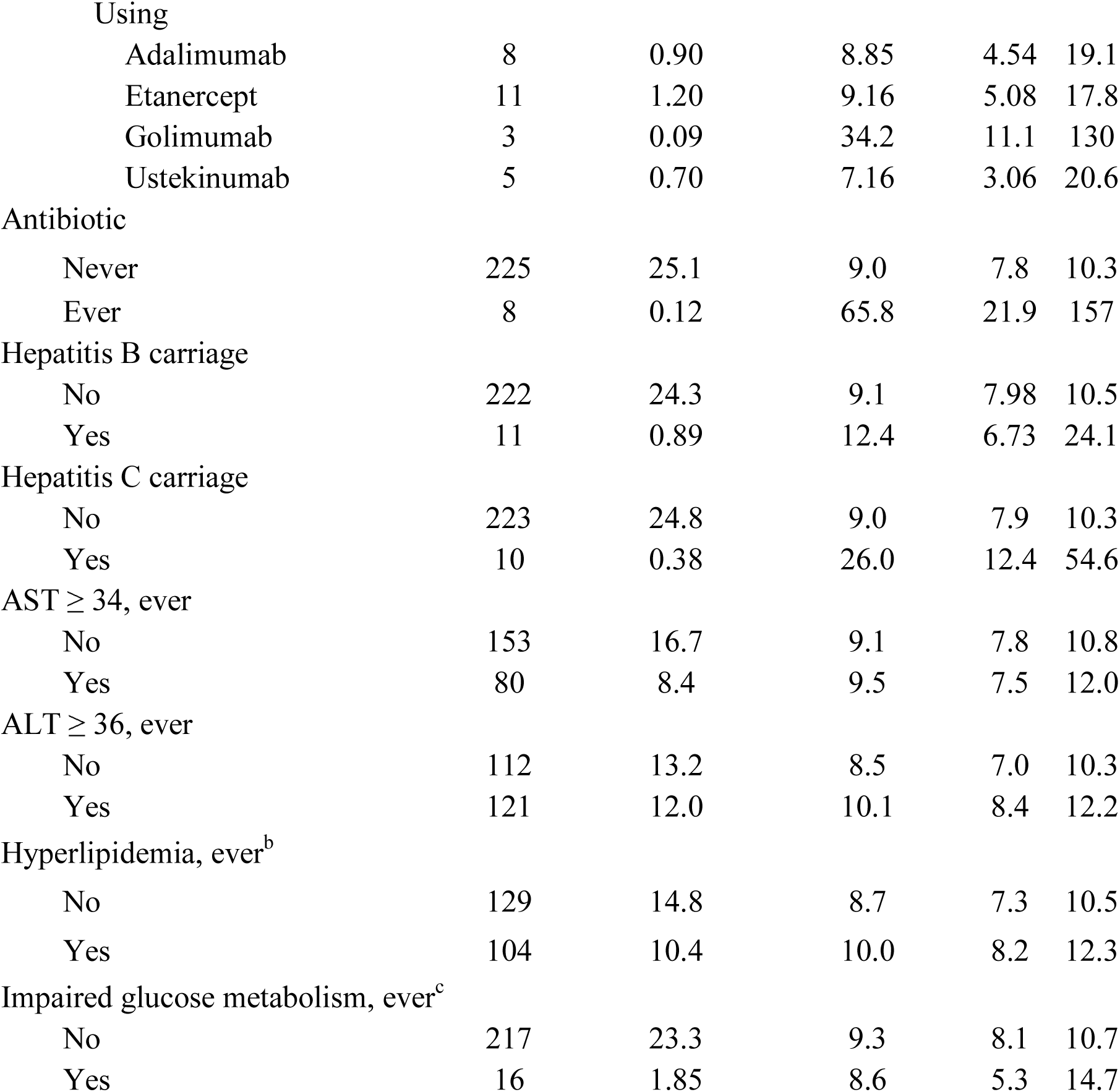
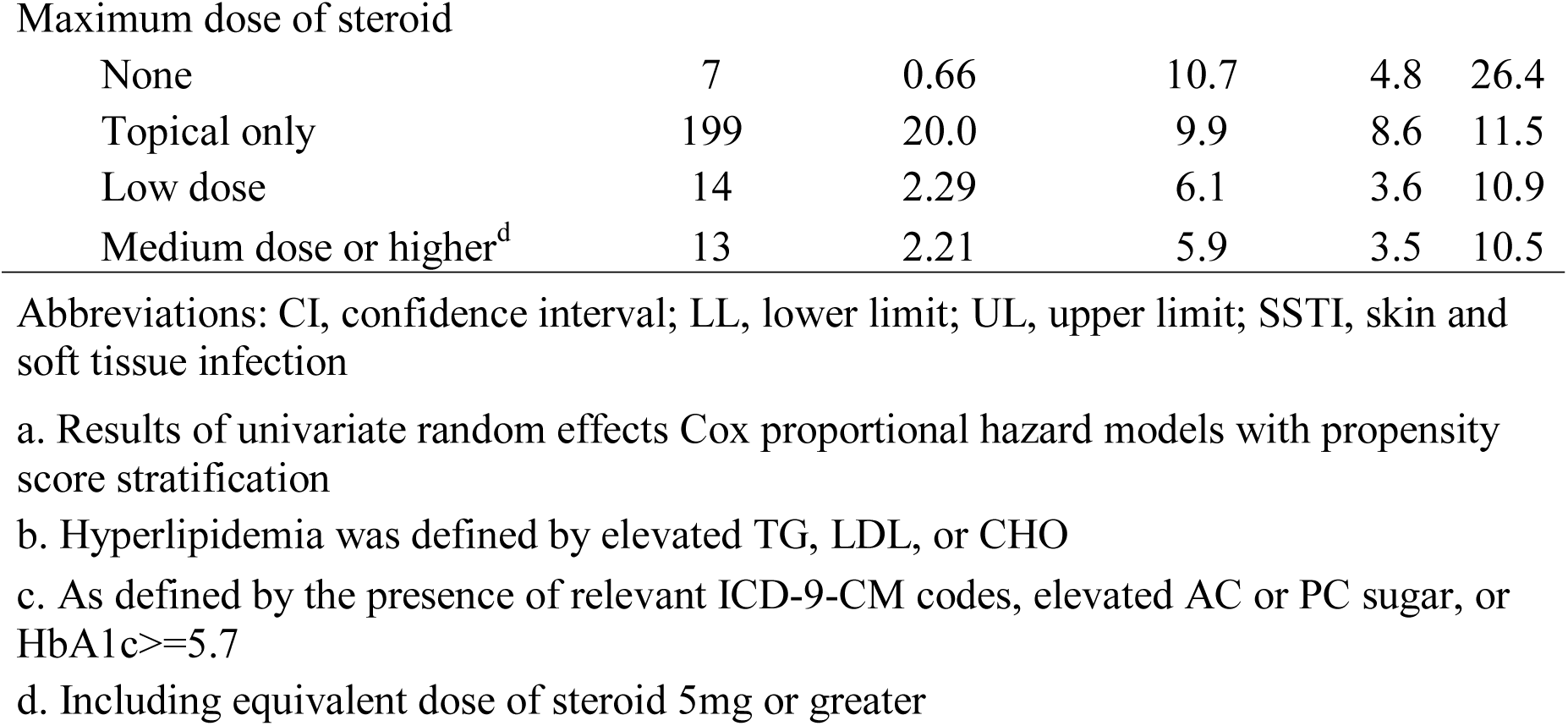
Incidence rates and crude hazard ratios of SSTI by selected characteristics among at-risk adult psoriasis patients in 2010-2015

| Characteristics | No.<br>events | Person-years<br>(in 100s) | Incidence rate<br>(per 100 py) | 95% CI |  |
| --- | --- | --- | --- | --- | --- |
|  |  |  |  | LL | UL |
| Total | 233 | 25.18 | 9.3 | 8.1 | 10.6 |
| Age, (year) |  |  |  |  |  |
| 20-40 | 55 | 6.07 | 9.1 | 6.9 | 12.0 |
| 40-65 | 139 | 15.6 | 8.9 | 7.5 | 10.6 |
| ≥ 65 | 39 | 3.47 | 11.2 | 8.2 | 15.7 |
| Sex |  |  |  |  |  |
| Female | 64 | 7.67 | 8.3 | 6.5 | 10.8 |
| Male | 169 | 17.5 | 9.6 | 8.2 | 11.3 |
| Calendar time at study entry |  |  |  |  |  |
| 2010 | 138 | 15.71 | 8.8 | 7.4 | 10.5 |
| 2011 | 28 | 3.83 | 7.3 | 5.0 | 11.0 |
| 2012 | 28 | 3.19 | 8.8 | 6.1 | 12.9 |
| 2013 | 30 | 1.95 | 15.4 | 10.7 | 22.8 |
| 2014 | 9 | 0.51 | 17.7 | 9.1 | 38.0 |
| Arthropathy, ever |  |  |  |  |  |
| No | 147 | 15.8 | 9.3 | 7.9 | 11.1 |
| Yes | 86 | 9.41 | 9.1 | 7.3 | 11.5 |
| Biologics use |  |  |  |  |  |
| Never | 200 | 19.2 | 10.4 | 9.0 | 12.1 |
| Ever | 33 | 6.0 | 5.5 | 4.0 | 7.9 |
| Not using | 6 | 3.09 | 1.94 | 0.89 | 5.1 |
| Using |  |  |  |  |  |
| Adalimumab | 8 | 0.90 | 8.85 | 4.54 | 19.1 |
| Etanercept | 11 | 1.20 | 9.16 | 5.08 | 17.8 |
| Golimumab | 3 | 0.09 | 34.2 | 11.1 | 130 |
| Ustekinumab | 5 | 0.70 | 7.16 | 3.06 | 20.6 |
| Antibiotic |  |  |  |  |  |
| Never | 225 | 25.1 | 9.0 | 7.8 | 10.3 |
| Ever | 8 | 0.12 | 65.8 | 21.9 | 157 |
| Hepatitis B carriage |  |  |  |  |  |
| No | 222 | 24.3 | 9.1 | 7.98 | 10.5 |
| Yes | 11 | 0.89 | 12.4 | 6.73 | 24.1 |
| Hepatitis C carriage |  |  |  |  |  |
| No | 223 | 24.8 | 9.0 | 7.9 | 10.3 |
| Yes | 10 | 0.38 | 26.0 | 12.4 | 54.6 |
| AST $\geq 34$, ever | | | | | |
| No | 153 | 16.7 | 9.1 | 7.8 | 10.8 |
| Yes | 80 | 8.4 | 9.5 | 7.5 | 12.0 |
| ALT $\geq 36$, ever | | | | | |
| No | 112 | 13.2 | 8.5 | 7.0 | 10.3 |
| Yes | 121 | 12.0 | 10.1 | 8.4 | 12.2 |
| Hyperlipidemia, ever<sup>b</sup> |  |  |  |  |  |
| No | 129 | 14.8 | 8.7 | 7.3 | 10.5 |
| Yes | 104 | 10.4 | 10.0 | 8.2 | 12.3 |
| Impaired glucose metabolism, ever<sup>c</sup> |  |  |  |  |  |
| No | 217 | 23.3 | 9.3 | 8.1 | 10.7 |
| Yes | 16 | 1.85 | 8.6 | 5.3 | 14.7 |
| Maximum dose of steroid |  |  |  |  |  |
| None | 7 | 0.66 | 10.7 | 4.8 | 26.4 |
| Topical only | 199 | 20.0 | 9.9 | 8.6 | 11.5 |
| Low dose | 14 | 2.29 | 6.1 | 3.6 | 10.9 |
| Medium dose or higher<sup>d</sup> | 13 | 2.21 | 5.9 | 3.5 | 10.5 |

Abbreviations: CI, confidence interval; LL, lower limit; UL, upper limit; SSTI, skin and soft tissue infection
- a. Results of univariate random effects Cox proportional hazard models with propensity score stratification - b. Hyperlipidemia was defined by elevated TG, LDL, or CHO - c. As defined by the presence of relevant ICD-9-CM codes, elevated AC or PC sugar, or HbA1c ≥ 5.7 - d. Including equivalent dose of steroid 5mg or greater

**Figure 2.**
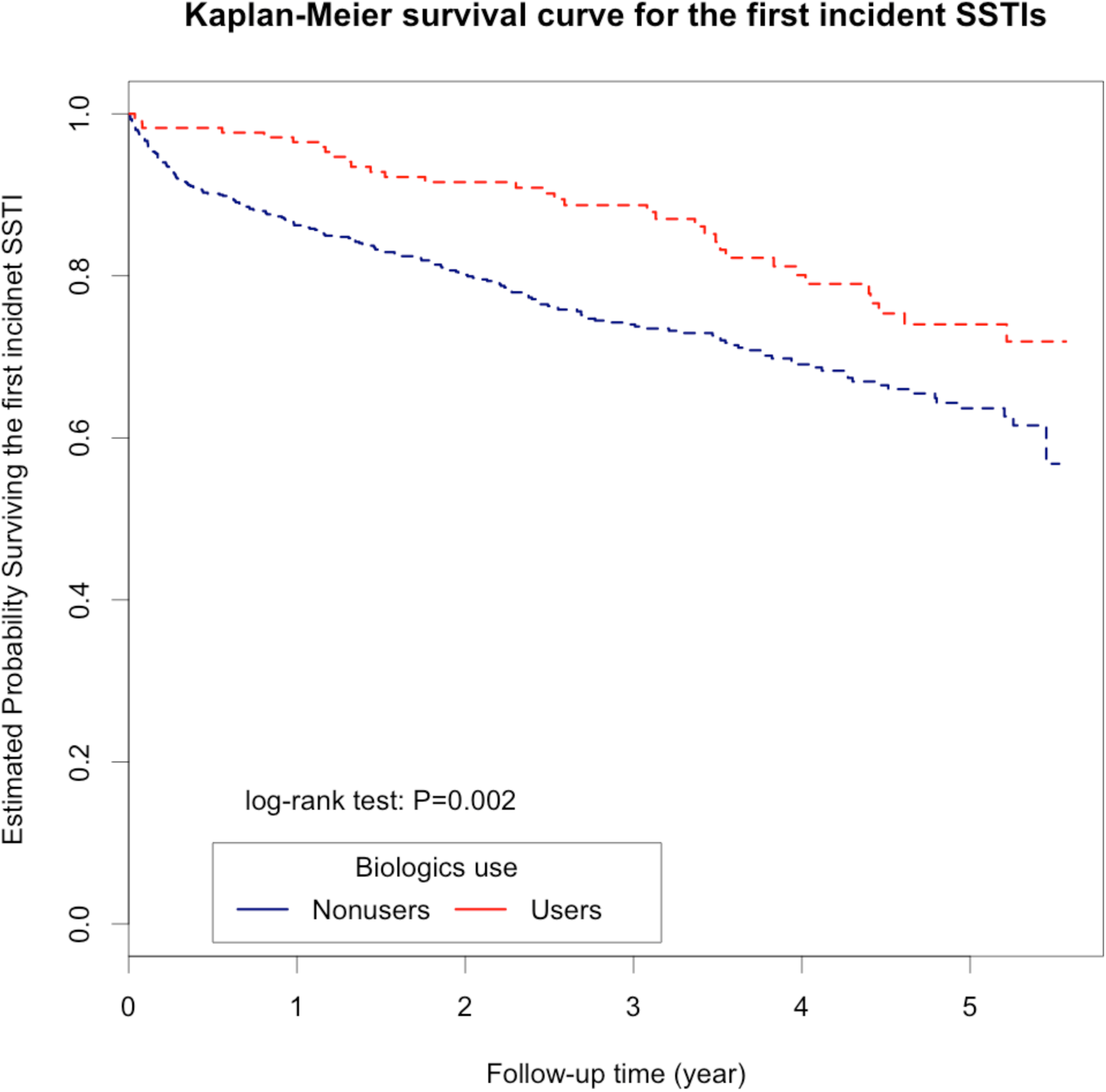
Kaplan-Meier survival curves for incident SSTIs for biologics users and nonusers, before propensity-score adjustment.

According to the estimated cumulative probabilities for incident SSTIs in patients by the strata of propensity scores, patients showed distinct SSTI risks based on their initial clinical characteristics alone (*P* for log-rank test: 0.005; Figure S1). Specifically, patients who would have been most likely to receive biologics (in quintile 5) had an 89% lower risk for SSTI than those with the lowest likelihood of using biologics (in quintile 1, crude HR: 0.11, 95%CI: 0.03, 0.41).

In general, there was an 89% risk reduction in biologics users versus nonusers (crude hazard ratio [HR]: 0.11, 95%CI: 0.05, 0.26; Table 3). For each 10-year increase in age, patients’ risks for SSTI decreased by 64% (crude HR: 0.36, 95%CI: 0.20, 0.65) whereas a recent entry into the study was associated with a 1.2-fold increase in SSTI risks (crude HR: 2.21, 95%CI: 1.56, 3.14). Most biologic agents prescribed were not correlated with an increased SSTI risk; notably, Ustekinumab was associated with an 81%-reduced risk as compared to nonbiologics (crude HR: 0.19, 95%CI: 0.05, 0.80). With additional adjustment for age, calendar time at entry, liver function, hyperlipidemia, and number of steroids in use, biologics users had a 74% lower SSTI risk than nonusers (adjusted HR: 0.26, 95%CI: 0.12, 0.58). When comparing time periods exposed to biologics with those exposed to nonbiologics in the subgroup of biologics users only, the former had a 21% higher risk for SSTIs than the latter (adjusted HR: 1.21, 95%CI: 0.41, 3.59) yet the association was not statistically significant (*P*: 0.740).

**Table 3.**
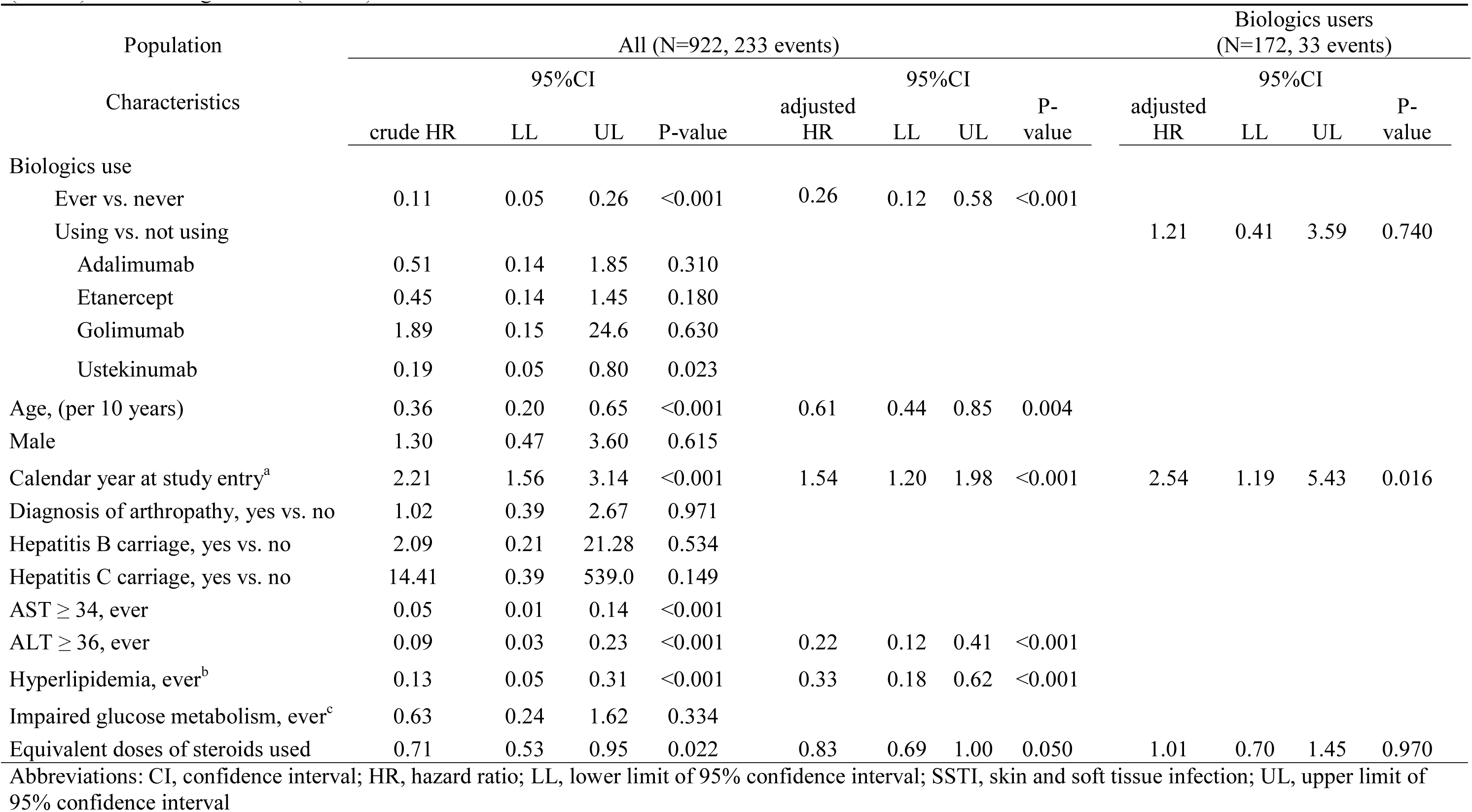
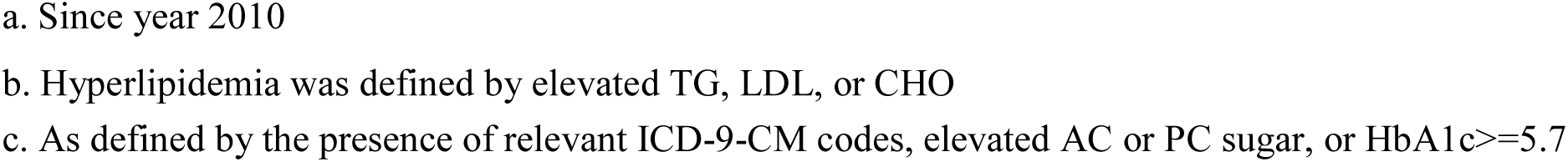
Results of propensity score-stratified, random effects Cox regression models comparing risks for incident SSTIs in at-risk adult psoriasis patients (N=922) and in biologics users (N=172)

## DISCUSSION

In a retrospective cohort of adult psoriasis patients, we did not find evidence for an increased SSTI risk associated with biologics use. When comparing time-periods spent on nonbiologics, biologics-ever users showed a comparable SSTI risk in time-periods exposed to biologics. In fact, as compared to nonusers, users showed a substantially low SSTI risk over the study period. Our finding was in contrary to previous reports similarly using retrospective clinical data.[15, 19] In reviewing medical charts of 398 Canadian patients in 2005-2014, Kim et al. found that incidence rates of infections leading to treatment cessation were comparably low (< 1/100 patient-years) for the biologics under study; yet there were no nonbiologics-associated infection rates reported.[15] Hadda and colleagues found that, biologics were associated with a higher overall risk for the first infection episode in psoriatic patients with arthritis (HR: 1.61, P: 0.001) but not in those without joint involvement (HR: 1.32, P: 0.73).[19]

Intuitively, the dominance of type I helper T cell (T_H_1) reactions seen in psoriatic tissues and in animal models[40] may appear paradoxical to reported patients’ susceptibility to bacterial colonization by observational studies.[5] In murine models, scientists have demonstrated that components of bacterial cell wall could induce hosts’ anti-inflammatory responses by down-regulating the production of pro-inflammatory cytokines including tumour necrosis factor-α and interferon-γ; antimicrobial peptides; and T cell activation.[41, 42] In vitro studies also showed that *S. aureus* could drive the clonal expansion of hosts’ T cells away from the pro-inflammatory, T_H_1/ T_H_17 pathway towards the anti-inflammatory regulatory T cell (T_reg_) pathway.[43, 44] Together, these molecular mechanisms of host-pathogen interactions may suggest an increased risk for bacterial colonization, and a potential (collateral) benefit of reduced colonization while blocking the pro-inflammatory pathway by biologics. Furthermore, despite of theoretical risks for increased susceptibility to intracellular pathogens, studies of interleukin 12/23p40 deficiencies suggested that both cytokines could be in redundant immune pathways against a number of human pathogens,[45] which was consistent with the generally low incidence of infections including SSTIs associated with biologics use in the trial setting.[7-10]

Besides a potential effectiveness in lowering SSTI risks by biologics, it was equally likely that the observed protective effects were biased results due to treatment selection as suggested by the literature.[22] As shown in Figure 3, comparisons between users and nonusers could be misleading if not accounting for the differential, pre-treatment susceptibility for SSTIs by group membership in a nonrandomized setting. Previously, Kalb et al. reported that there was a 2-fold risk for cellulitis, abscess and (other) skin infections in the biologics group as compared to the nonbiologics/ non-methotrexate group;[22] however, the authors did not provide adjusted estimates despite that patients using biologics were apparently more obese than their counterparts and a well-studied link of obesity itself to an increased SSTI risk.[26, 46-48]

Conventional regression models may yield extrapolation-based results in the presence of multiple, imbalanced prognostic factors across comparators as is frequently encountered in the clinical practice.[27, 49] The current analysis sought to apply propensity score stratification to create balanced distributions of important factors across the propensity-score strata (Figures S1-S2). Nevertheless, a strong negative association between patients’ age and SSTI risks (Table 2, Figure S3) indicated the presence of residual confounding by indication. In other words, older patients were much less likely to receiving biologics than younger ones (Table 1), thus a lower biologics-associated SSTI risk; yet the small sample size of users prohibited a formal interaction test by age on the biologics-SSTI relationship. Likewise, the seemingly protective effects of an elevated serum level of ALT and history of hyperlipidemia outlined the highly selective nature of the patient group eligible for biologics therapeutics as indicated by either clinical guidelines[ 26, 50, 51] or the national insurance reimbursement policy in Taiwan,[52] or both.

There are limitations in the current analysis worth reminding before generalizing the study findings. To tackle the incomparability issue inherent to observational data, the propensity score method appeared only partially effective. The lack of detailed disease history, previous use of biologics, and history of systemic infections could result in misclassifying individuals into strata of a lower (or under-estimated) probability to receive biologics. Such misclassification bias might contribute to the between-group incomparability, leading towards the residual confounding by indication. Notably, the lack of disease severity scores or quality of life indices prior to treatment, which were important determinants for prescribing biologics, did not seem to correlate with infection risks in psoriasis patients.[10, 19, 22]

Second, assuming person-time equivalence, we also determined biologics’ effect on SSTI risks only among biologics users, among whom we also assumed exchangeability between the biologics-exposed and the yet unexposed users.[49] This assumed comparability among users should be optimally aligned by disease duration, the information of which was lacking in the current analysis. The small number of SSTI events in the subgroup analysis might also render the comparisons nonsignificant.

## Conclusions

We found no evidence that biologics use was associated with increased SSTI risks in adult psoriasis patients either in between-group or in within-group comparisons. Given pragmatic constraints, future comparative studies on biologics-associated health effects shall include only biologics users and perform cross-over comparisons so as to ensure comparability and valid inferences.

## List of abbreviations

S. aureus: Staphylococcus aureus
SSTI: skin and soft tissue infection
MRSA: methicillin-resistant S. aureus
AD: atopic dermatitis
ICD-9-CM: International Classification of Diseases, Ninth Revisions, Clinical Modification
EMD: Electronic Medical Database
AST: aspartate transaminase
ALT: alanine transaminase
LDL: low density lipoprotein-cholesterol
AIC: Akaike information criteria
IQR: interquartile range
IR: incidence rate
CI: confidence interval
HR: hazard ratio
T_H_1: type I helper T cell
T_reg_: regulatory T cell

## DECLARATIONS

### Ethics approval and consent to participate

The institutional review board at Chang Gung Memorial Hospital has reviewed and approved the study protocol and the analytic plan. The requirement for consents from participants was also waved as this was a retrospective analysis of existing data.

### Consent for publication

Not applicable.

### Availability of data and material

The datasets analysed in the current study are not publicly available due to the institutional regulations on individual medical information. However, de-identified secondary data and statistical codes are available from the corresponding author upon reasonable request.

### Funding

This work was supported by Chang Gung Medical Foundation [CMRPG3E1971, CMRPG3E1972 to SHL]. However, the funder had no role in study design, data collection, data analysis, manuscript preparation or publication decisions.

### Authors’ contributions

SL conceived and designed the study, collected the data, performed statistical analysis, interpreted results of analysis, and drafted the manuscript. TH and LC helped clean the original data, performed statistical analysis, and contributed to data interpretation and manuscript drafting. YCH and YHH participated in the study design, data collection, results interpretation, and preparation of the manuscript. All authors have reviewed and approved the submitted version of the manuscript.

### Competing interests

The authors declared that they have no competing interests.

## REFERENCES

1. Miller LS, Cho JS: Immunity against Staphylococcus aureus cutaneous infections. Nat Rev Immunol 2011, 11(8): 505–518.

2. Wertheim HFL, Melles DC, Vos MC, Van Leeuwen W, Van Belkum A, Verbrugh HA, Nouwen JL: The role of nasal carriage in Staphylococcus aureus infections. Lancet Infect Dis 2005, 5: 751–762.

3. Chuang YY, Huang YC: Molecular epidemiology of community-associated meticillin-resistant Staphylococcus aureus in Asia. Lancet Infect Dis 2013, 13(8): 698–708.

4. Boguniewicz M, Leung DY: Atopic dermatitis: a disease of altered skin barrier and immune dysregulation. Immunol Rev 2011, 242(1): 233–246.

5. Tomi NS, Kranke B, Aberer E: Staphylococcal toxins in patients with psoriasis, atopic dermatitis, and erythroderma, and in healthy control subjects. J Am Acad Dermatol 2005, 53(1): 67–72.

6. Albrich WC, Harbarth S: Health-care workers: source, vector, or victim of MRSA? Lancet Infect Dis 2008, 8(5): 289–301.

7. Gordon KB, Papp KA, Langley RG, Ho V, Kimball AB, Guzzo C, Yeilding N, Szapary PO, Fakharzadeh S, Li S et al: Long-term safety experience of ustekinumab in patients with moderate to severe psoriasis (Part II of II): results from analyses of infections and malignancy from pooled phase II and III clinical trials. J Am Acad Dermatol 2012, 66(5): 742–751.

8. Gordon KB, Blauvelt A, Papp KA, Langley RG, Luger T, Ohtsuki M, Reich K, Amato D, Ball SG, Braun DK et al: Phase 3 Trials of Ixekizumab in Moderate-to-Severe Plaque Psoriasis. N Engl J Med 2016, 375(4): 345–356.

9. Papp KA, Griffiths CE, Gordon K, Lebwohl M, Szapary PO, Wasfi Y, Chan D, Hsu MC, Ho V, Ghislain PD et al: Long-term safety of ustekinumab in patients with moderate-to-severe psoriasis: final results from 5 years of follow-up. Br J Dermatol 2013, 168(4): 844–854.

10. Carretero G, Ferrandiz C, Dauden E, Vanaclocha Sebastian F, Gomez-Garcia FJ, Herrera-Ceballos E, De la Cueva-Dobao P, Belinchon I, Sanchez-Carazo JL, Alsina-Gibert M et al: Risk of adverse events in psoriasis patients receiving classic systemic drugs and biologics in a 5-year observational study of clinical practice: 2008-2013 results of the Biobadaderm registry. J Eur Acad Dermatol Venereol 2015, 29(1): 156–163.

11. Henseler T, Christophers E: Disease concomitance in psoriasis. J Am Aca Dermatol 1995, 32: 982–986.

12. Fry L, Baker BS: Triggering psoriasis: the role of infections and medications. Clin Dermatol 2007, 25(6): 606–615.

13. Balci DD, Duran N, Ozer B, Gunesacar R, Onlen Y, Yenin JZ: High prevalence of Staphylococcus aureus cultivation and superantigen production in patients with psoriasis. Eur J Dermatol 2009, 19(3): 238–242.

14. Ng CY, Huang YH, Chu CF, Wu TC, Liu SH: Risks for Staphylococcus aureus Colonization in Psoriasis Patients: A Systematic Review and Meta-Analysis. (In press) 2017.

15. Kim WB, Marinas JE, Qiang J, Shahbaz A, Greaves S, Yeung J: Adverse events resulting in withdrawal of biologic therapy for psoriasis in real-world clinical practice: A Canadian multicenter retrospective study. J Am Acad Dermatol 2015, 73(2): 237–241.

16. Papp K, Gottlieb AB, Naldi L, Pariser D, Ho V, Goyal K, Fakharzadeh S, Chevrier M, Calabro S, Langholff W et al: Safety Surveillance for Ustekinumab and Other Psoriasis Treatments From the Psoriasis Longitudinal Assessment and Registry (PSOLAR). Journal of drugs in dermatology : JDD 2015, 14(7): 706–714.

17. Pereira R, Lago P, Faria R, Torres T: Safety of Anti-TNF Therapies in Immune-Mediated Inflammatory Diseases: Focus on Infections and Malignancy. Drug Dev Res 2015, 76(8): 419–427.

18. Sorenson E, Koo J: Evidence-based adverse effects of biologic agents in the treatment of moderate-to-severe psoriasis: Providing clarity to an opaque topic. The Journal of dermatological treatment 2015, 26(6): 493–501.

19. Haddad A, Li S, Thavaneswaran A, Cook RJ, Chandran V, Gladman DD: The Incidence and Predictors of Infection in Psoriasis and Psoriatic Arthritis: Results from Longitudinal Observational Cohorts. J Rheumatol 2016, 43(2): 362–366.

20. Gniadecki R, Bang B, Bryld LE, Iversen L, Lasthein S, Skov L: Comparison of long-term drug survival and safety of biologic agents in patients with psoriasis vulgaris. Br J Dermatol 2015, 172(1): 244–252.

21. van den Reek JM, Tummers M, Zweegers J, Seyger MM, van Lumig PP, Driessen RJ, van de Kerkhof PC, Kievit W, de Jong EM: Predictors of adalimumab drug survival in psoriasis differ by reason for discontinuation: long-term results from the Bio-CAPTURE registry. J Eur Acad Dermatol Venereol 2015, 29(3): 560–565.

22. Kalb RE, Fiorentino DF, Lebwohl MG, Toole J, Poulin Y, Cohen AD, Goyal K, Fakharzadeh S, Calabro S, Chevrier M et al: Risk of Serious Infection With Biologic and Systemic Treatment of Psoriasis: Results From the Psoriasis Longitudinal Assessment and Registry (PSOLAR). JAMA Dermatol 2015, 151(9): 961–969.

23. Grijalva CG, Chen L, Delzell E, Baddley JW, Beukelman T, Winthrop KL, Griffin MR, Herrinton LJ, Liu L, Ouellet-Hellstrom R et al: Initiation of tumor necrosis factor-alpha antagonists and the risk of hospitalization for infection in patients with autoimmune diseases. JAMA 2011, 306(21): 2331–2339.

24. Shalom G, Zisman D, Bitterman H, Harman-Boehm I, Greenberg-Dotan S, Dreiher J, Feldhamer I, Moser H, Hammerman A, Cohen Y et al: Systemic Therapy for Psoriasis and the Risk of Herpes Zoster: A 500,000 Person-year Study. JAMA Dermatol 2015, 151(5): 533–538.

25. Menter A, Gottlieb A, Feldman SR, Van Voorhees AS, Leonardi CL, Gordon KB, Lebwohl M, Koo JYM, Elmets CA, Korman NJ et al: Guidelines of care for the management of psoriasis and psoriatic arthritis-Section 1. Overview of psoriaisis and guidelines of care for the treatment of psoriasis with biologics. J Am Acad Dermatol 2008, 58: 826–850.

26. Menter A, Korman NJ, Elmets CA, Feldman SR, Gelfand JM, Gordon KB, Gottlieb A, Koo JY, Lebwohl M, Leonardi CL et al: Guidelines of care for the management of psoriasis and psoriatic arthritis: section 6. Guidelines of care for the treatment of psoriasis and psoriatic arthritis: case-based presentations and evidence-based conclusions. J Am Acad Dermatol 2011, 65(1): 137–174.

27. Biondi-Zoccai G, Romagnoli E, Agostoni P, Capodanno D, Castagno D, D’Ascenzo F, Sangiorgi G, Modena MG: Are propensity scores really superior to standard multivariable analysis? Contemp Clin Trials 2011, 32(5): 731–740.

28. Rosenbaum PR, Rubin DB: The central role of the propensity score in observational studies for causal effects. Biometrika 1983, 70(1): 41–55.

29. Schneeweiss S, Rassen JA, Glynn RJ, Avorn J, Mogun H, Brookhart MA: Highdimensional Propensity Score Adjustment in Studies of Treatment Effects Using Health Care Claims Data. Epidemiology 2009, 20: 512–522.

30. Vittinghoff E, Glidden DV, Shiboski SC, McCulloch CE: Regression Methods in Biostatistics: Linear, Logistic, Survival, and Repeated Measures Models. New York, USA: Springer Science+Business Media, LLC; 2012.

31. Suaya JA, Eisenberg DF, Fang C, Miller LG: Skin and soft tissue infections and associated complications among commercially insured patients aged 0-64 years with and without diabetes in the U.S. PLoS One 2013, 8(4).

32. Clinical Calculators: Steroid Equivalence Converter [http://www.medcalc.com/steroid.html]

33. Therneau TM, Grambsch PM: Modeling survival data, extending the Cox model, vol. 1st. New York: Springer-Verlag; 2000.

34. Hosmer DW, Lemeshow S, May S: Applied Survival Analysis: Regression Modeling of Time-to-Event Data, 2nd edn. Hoboken, New Jersey: John Wiley & Sons, Inc.; 2008.

35. Greenland S: Introduction to Regression Modeling. In: Modern Epidemiology. edn. Edited by Rothman KJ, Greenland S, Lash TL. Philadelphia, PA 19106 USA: Lippincott Williams & Wilkins; 2008: 412–455.

36. Austin PC: The use of propensity score methods with survival or time-to-event outcomes: reporitng measures of effect similar to those used in randomized experiments. Stat Med 2014, 33: 1242–1258.

37. StataCorp LP: Stata Statistical Software: Release 13. In., 13.1 edn. College Station, TX; 2013.

38. R Core Team. R: A language and environment for statistical computing. In., 3.3.1 edn. Vienna, Austria: R Foundation for Statistical Computing; 2016.

39. Therneau T, Lumley T: Package ‘survival’: Survival Analysis. In., 2.39.5 edn; 2016: Contains the core survival analysis routines, including definition of Surv objects, Kaplan-Meier and Aalen-Johansen (multi-state) curves, Cox models, and parametric accelerated failure time models.

40. Nestle FO, Di Meglio P, Qin JZ, Nickoloff BJ: Skin immune sentinels in health and disease. Nat Rev Immunol 2009, 9(10): 679–691.

41. Chau TA, McCully ML, Brintnell W, An G, Kasper KJ, Vines ED, Kubes P, Haeryfar SM, McCormick JK, Cairns E et al: Toll-like receptor 2 ligands on the staphylococcal cell wall downregulate superantigen-induced T cell activation and prevent toxic shock syndrome. Nat Med 2009, 15(6): 641–648.

42. Lai Y, Di Nardo A, Nakatsuji T, Leichtle A, Yang Y, Cogen AL, Wu ZR, Hooper LV, Schmidt RR, von Aulock S et al: Commensal bacteria regulate Toll-like receptor 3-dependent inflammation after skin injury. Nat Med 2009, 15(12): 1377–1382.

43. Rabe H, Nordstrom I, Andersson K, Lundell AC, Rudin A: Staphylococcus aureus convert neonatal conventional CD4(+) T cells into FOXP3(+) CD25(+) CD127(low) T cells via the PD-1/PD-L1 axis. Immunology 2014, 141(3): 467–481.

44. Zielinski CE, Mele F, Aschenbrenner D, Jarrossay D, Ronchi F, Gattorno M, Monticelli S, Lanzavecchia A, Sallusto F: Pathogen-induced human TH17 cells produce IFN-gamma or IL-10 and are regulated by IL-1beta. Nature 2012, 484(7395): 514–518.

45. Teng MW, Bowman EP, McElwee JJ, Smyth MJ, Casanova JL, Cooper AM, Cua DJ: IL-12 and IL-23 cytokines: from discovery to targeted therapies for immune-mediated inflammatory diseases. Nat Med 2015, 21(7): 719–729.

46. Huttunen R, Syrjanen J: Obesity and the risk and outcome of infection. Int J Obes (Lond) 2013, 37(3): 333–340.

47. Phung DT, Wang Z, Rutherford S, Huang C, Chu C: Body mass index and risk of pneumonia: a systematic review and meta-analysis. Obes Rev 2013, 14(10): 839–857.

48. Tateya S, Kim F, Tamori Y: Recent advances in obesity-induced inflammation and insulin resistance. Front Endocrinol (Lausanne) 2013, 4: 93.

49. Little RJ, Rubin DB: Causal effects in clinical and epidemiological studies via potential outcomes: concepts and analytical approaches. Annu Rev Public Health 2000, 21: 121–145.

50. Ohtsuki M, Terui T, Ozawa A, Morita A, Sano S, Takahashi H, Komine M, Etoh T, Igarashi A, Torii H et al: Japanese guidance for use of biologics for psoriasis (the 2013 version). J Dermatol 2013, 40(9): 683–695.

51. Gossec L, Smolen JS, Ramiro S, de Wit M, Cutolo M, Dougados M, Emery P, Landewe R, Oliver S, Aletaha D et al: European League Against Rheumatism (EULAR) recommendations for the management of psoriatic arthritis with pharmacological therapies: 2015 update. Ann Rheum Dis 2016, 75(3): 499–510.

52. The National Health Insurance Pharmaceutical Benefits and Reimbursement Schedule: The reimbursement restrictions of pharmaceuticals. In. Taipei, Taiwan: Ministry of Health and Welfare.

